# Explorations with Conditioned Stimuli in Autoshaping Procedures

**DOI:** 10.1101/475541

**Authors:** Jill F. Nehrbas, Elizabeth B. Smedley, S. Smith Kyle

## Abstract

Sign-tracking is a form of autoshaping where by animals reliably develop conditioned responses toward stimuli that predict an outcome. While the assignment of some value to a predictive cue may be adaptive (i.e., to be alerted to food and water sources), the attribution of value to predictive cues can be maladaptive as seen in behaviors elicited during addiction. Here we test if responding to the predictive cue changes in the context of other cues that are only partially predictive (Experiment 1). Previous work on sequential cues leading to reward have shown a bias in responding toward the first cue in the sequence over learning (Smedley and Smith 2018a, 2018b). Here we test if this effect is unique to discrete cues or if a bias in responding can be seen in a single, long cue (Experiment 2). Finally, we investigate if sign-tracking responses can reliably develop towards a cue that arrives after the delivery of reward (backwards conditioning, Experiment 3). Together, we aim to address various gaps in knowledge about the nature of the sign-tracking response.

## Introduction

When a cue reliably predicts the presentation of a reward such as food, animals may attribute motivational value to the cue itself. Autoshaping or sign-tracking behaviors involve the animal interacting with a discrete and localized conditioned stimulus, despite receiving the reward regardless of their actions. While sign-trackers may benefit by using signals in their environment to guide them to food and water necessary for survival, it is possible that the attribution of incentive salience to cues could lead to negative motivational behaviors such as addiction (Flagel and Robinson, 2017). Research has shown that unlike goal-trackers, who more frequently go directly to where the reward is delivered, sign-trackers attribute more incentive salience to cues preceding drug-intake, and these cues can be more likely to trigger drug addiction relapse (Berridge and Robinson, 2016).

Past research has investigated how partially reinforced sign-tracking cues affect sign-tracking behavior and found that partial reinforcement of the conditioned stimulus resulted in more sign-tracking behavior while full reinforcement led to more frequent goal-tracking behavior. They also found that responding declined more slowly to the partially reinforced stimulus compared to the fully reinforced stimulus during extinction (Davey and Cleland, 1982). Additionally, research into the effects of serially occurring cues has shown that animals favor responding to the reward distal cue over the reward proximal cue and, therefore, that the incentive salience of one cue can influence the value of the other cue (Smedley and Smith 2018a, 2018b). Research, however, has not investigated how a partially reinforced cue could alter responding to a fully reinforced cue.

While there is limited literature that directly looks into the effects of varying the length of presentation of the conditioned stimulus, previous studies have shown that in a sequential presentation of lever cues the lever presented furthest in time from the reward delivery acquires more lever responding than the lever temporally closer in time to the reward over sessions (Smedley and Smith 2018a, 2018b). It is important to investigate whether this phenomenon is unique to discrete lever presentations or if presenting a conditioned stimulus for a longer amount of time will result in similar responding over sessions.

Sign-tracking is almost always conducted with a predictive cue that comes prior to reward delivery and earlier conditioning models such as Wagner’s SOP Model assume that the CS must precede the US for a predictive relationship to be formed (Rescorla & Wagner, 1972). However, research has suggested that the backward CS can become an inhibitor, signaling the start of the intertrial period (Chang, Blaisdell, & Miller, 2003; Maier, Rapaport, & Wheatley, 1976; Moscovitch & LoLordo, 1968; Plotkin & Oakley, 1975; Siegel & Domjan, 1971, 1974). It remains unclear how animals might attribute value to a cue that always follows the delivery of the reward.

Here we address if manipulation of the reward structure of one lever cue impacts the responding towards a fully reinforced lever (Experiment 1). We investigate if the serial effect seen previously (Smedley and Smith 2018a, 2018b) is related to discrete cues or if the length of the cue matters. We test if pressing occurs earlier in the lever presentation as sessions progress (Experiment 2). Finally, we address if sign-tracking occurs in a backwards conditioning paradigm (Experiment 3). Together, these results will clarify uncertainties about the nature of the sign-tracking response.

## Materials and Methods

### Animals

Long Evans rats obtained from Charles River (n=40; Charles River, Indianapolis, IN, USA) were single housed in ventilated plastic cages in a climate-controlled colony room on a 12 h light/dark cycle (lights on at 7:00 A.M.). Experiments were conducted during the light cycle. Weight was restricted to 85% ad libitum weight prior to magazine training and maintained throughout all experimental procedures. Rats were provided with standard rat chow (Harlan Teklad 2014) after each testing session to maintain 85% weight and were given free access to water the remaining 23 h/d. All procedures were approved by the Dartmouth College Animal Care and Use Committee.

### Apparatus

Training and testing were conducted in standard operant chambers (Med Associates, St. Albans, VT), which were enclosed in sound- and light-attenuating cabinets outfitted with fans for ventilation and white noise. Chambers contained two retractable levers to the left and right of a recessed magazine. Lever deflections were recorded automatically. Magazine entries were recorded through breaks in an infrared beam adjacent to the magazine.

### Behavioral measures and analyses

Lever deflections, magazine entries, and time spent in magazine were recorded through MedPC. Presses per minute (ppm) are the total presses divided by the total available lever time divided by 60 minutes. Presses indicates a count of the number of downward deflections of the lever. Magazine entries reflects a count of the number of photobeam breaks into the magazine. Magazine entries per minute (mpm) is the total magazine entries divided by 60 minutes.

### Statistical Analyses

All statistical tests were carried out using R (R Core Team, 2013). Categorical variables with multiple levels were dummy coded to make predetermined comparisons between levels. All linear mixed models are fit by maximum likelihood and t-tests use Satterthwaite approximations of degrees of freedom (R; “lmerTest”, Kuznetsova et al. 2015). The reported statistics will include parameter estimates (β values), confidence intervals (95% bootstrapped confidence intervals around dependent variable), standard error of the parameter estimate (SE), and p-values (R; “lmerTest”, Kuznetsova et al. 2015). T-tests, Welch two sample t-tests for unpaired tests and paired t-tests, report t-values, 95% CIs, degrees of freedom, and p-values. Graphs were created through R (R Core Team, 2013) or GraphPad Prism (version 7.0a) and designed with Adobe Illustrator.

## Experiment 1: CS manipulation

Training began with a single, 35-min magazine acclimation session in which pellets were delivered with a probability of 1 pellet every 30 sec (approx. 60 pellets delivered over the session). Rats in Group Control (n=8) were then exposed to 12 daily, 60 min sessions of conditioning. In each session, rats received 25, 10 sec lever presentations (CS+) followed by lever retraction and reward delivery. Two pellets (BioServ, 45 mg Dustless Precision Pellets) were delivered followed by an average 60 sec ITI. The CS+ trials were pseudorandomly interspersed with 25 presentations of a 10 sec unreinforced lever (the CS-was the opposite lever in the chamber). Rats in Group Experimental (n=8) were also given 12 days of 60 min sessions, during which 25 presentations of a CS+ lever was always reinforced and was presented for 10 sec followed by reward delivery. The CS-lever was reinforced with a probability of 50%, such that on average every other trial was reinforced. When the CS-was reinforced, it was followed by the delivery of two pellets. Both groups then received four days of 50 trial extinction sessions (i.e., levers were presented and feeder noises occurred as during training but no pellets were given; Fig. 1A). All groups were counterbalanced with right or left levers as CS+ cues.

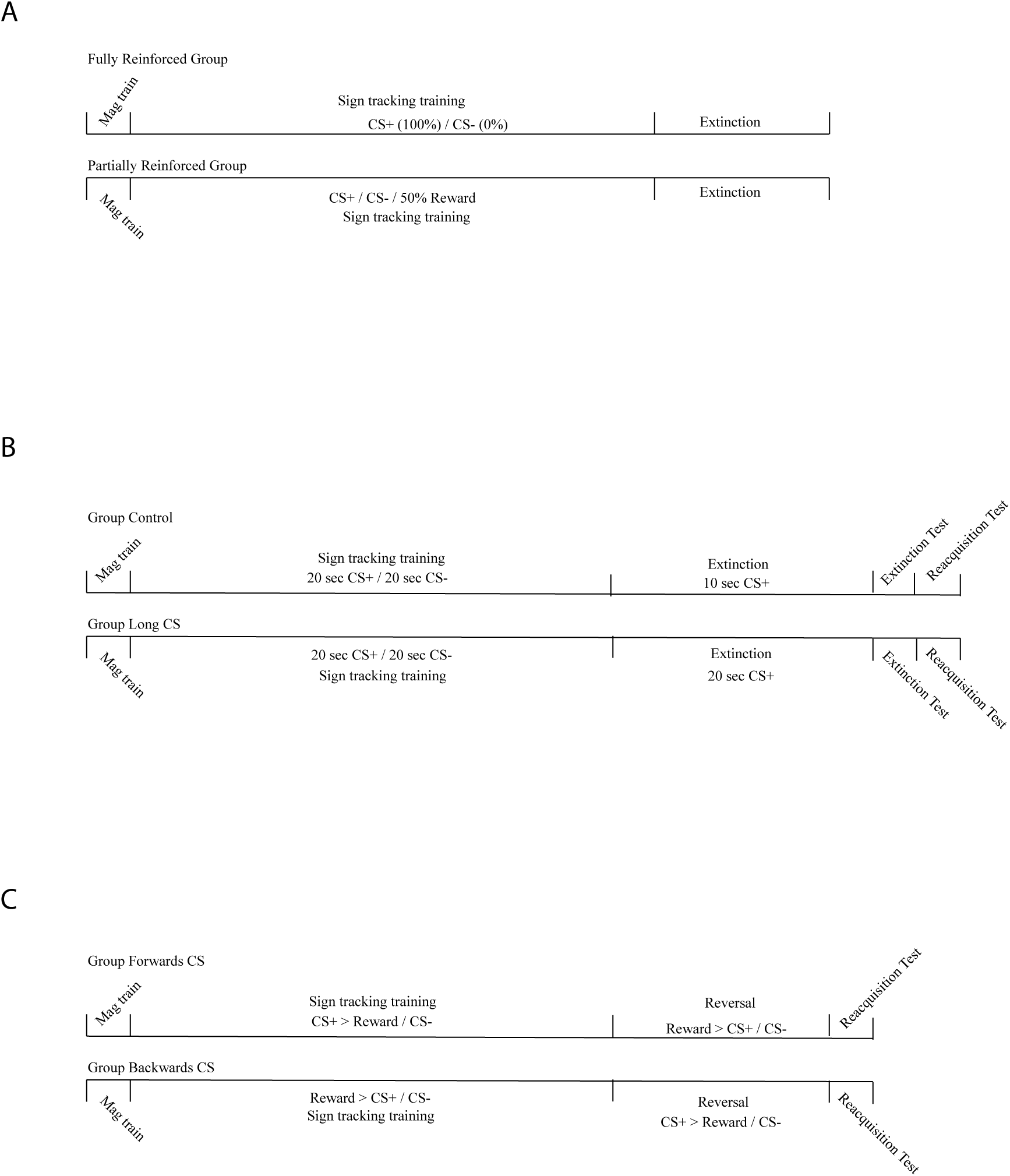

## Experiment 2: Long CS

Training began with a single, 35-min magazine acclimation session in which pellets were delivered with a probability of 1 pellet every 30 sec. Rats in Group Long CS (n=8) were then exposed to 12 daily, 60 min sessions of conditioning. In each session, rats received 25, 20 sec CS+ lever presentation followed by reward delivery of two pellets (BioServ, 45 mg Dustless Precision Pellets). The 25 CS+ trials were followed by an average 120 sec ITI. In addition to the CS+ trials, 25, 20 sec CS-trials occured throughout the session where the opposite lever in the chamber was delivered for 20 sec followed by an average 120 sec ITI. After 12 days of training, Group Long CS received five days of 50 trial extinction sessions (i.e., CS+ levers were presented and feeder noises occurred as during training but no pellets were given). Group Control continued with the same training paradigm they had received during the first 12 days, except no trials were rewarded. Extinction was followed by an extinction test (received training paradigm in extinction) and a reacquisition test (received training paradigm; Fig. 1B). All groups were counterbalanced with right or left levers as CS+ levers.

## Experiment 3: Backwards CS

Training began with a single, 35-min magazine acclimation session in which pellets were delivered with a probability of 1 pellet every 30 sec. Group Forward CS received 25 trials of a 10 sec presentation of a lever followed by the delivery of two pellets. An average 120 sec ITI separated the CS+ trials from the delivery of CS-trials (presentation of the opposite lever in the chamber followed by no reward delivery). Trial structure pseudorandomly delivered the CS+ or CS-trials, limiting to no more than two of either in sequence. Group Backward CS received 25 trials of two pellets being delivered (4 sec delivery period) followed by the 10 sec presentation of the CS+ lever. CS-trials were the delivery of the opposite lever in the chamber followed by nothing. Trial structure was the same as Group Forward CS (Fig. 1C).

## Results

### Experiment 1: CS Manipulation Training

#### Q1. Is CS+ (exp) different from CS+ (control)?

The presence of a differently reinforced lever did not impact presses toward the fully reinforced lever; all animals pressed similarly to a fully reinforced lever (Fig. 2A). A linear mixed model of press rates on the reinforced lever by group (exp v. control) by session and random intercepts (Eq.1; the addition of random slopes resulted in failed convergence of the model and was thus removed from the final model) revealed no significant main effect of group (Fig. 2A; est: −4.40 ppm; CI: −16.1-6.92; SE: 5.80; p = 0.454). Regardless of group, press rates did not differ over sessions (est: −0.15 ppm; CI: −0.85-0.45; SE: 0.32; p = 0.632). Groups did not differ over sessions (est: 0.41 ppm; CI: −0.48-1.31; SE: 0.457; p = 0.375).

**Figure 2.**
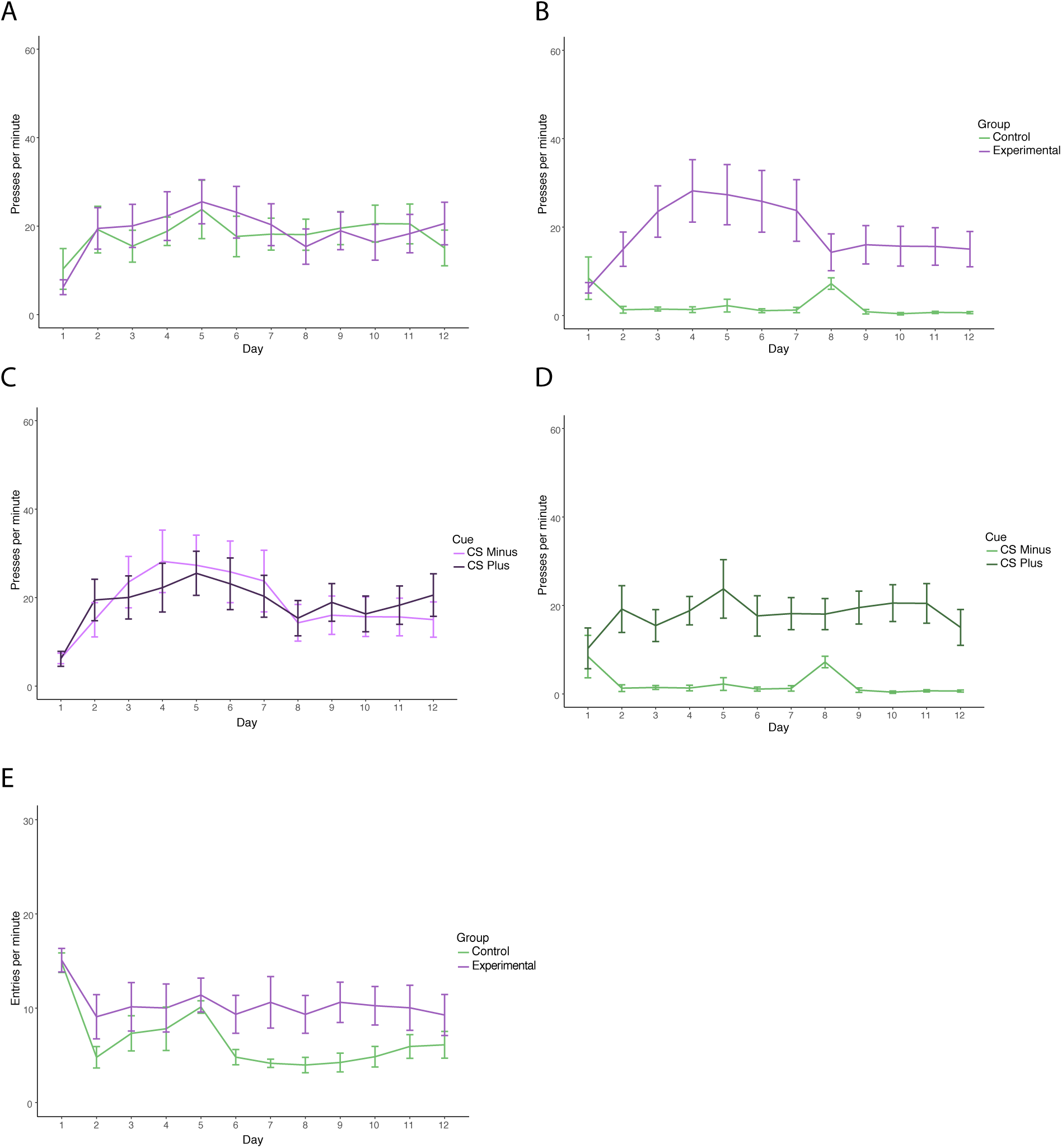
Experiment 1: Training. Training data from the first 12 days of sign-tracking in the CS manipulation experiment. Error bars are standard error of the mean. (A) Presses per minute on the CS+ by group (Control; 100 −0; Experimental; 100 −50). (B) Presses per minute on the CS-by group. Group 1 (Control; 100 −0) Group 2 (Experimental; 100 −50). (C) Presses per minute on CS+ vs. CS-(Experimental Group). CS-(50% partially reinforced; light pink) and CS+ (100% reinforced; dark purple). (D) Presses per minute on CS+ vs. CS-(Control Group). CS-(0% reinforced; light green) and CS+ (100% reinforced; dark green). (E) Magazine entries per minute by group over the 12 days of training (Control, 100 − 0, green; Experimental, 100 − 50, purple)

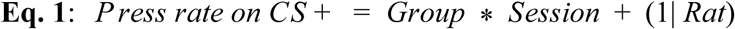

#### Q2. Is CS-(partially reinforced; exp) different from CS-(never reinforced; control)?

Animals pressed more toward a partially reinforced CS (group exp) than animals who received a never reinforced CS (group control; Fig. 2B). A linear mixed model of press rate by group (cont.V. exp) by session with random effects of slope and rat (Eq. 2) revealed a significant effect of group (Fig. 2B; est: 15.9 ppm; CI: 6.28-25.2; SE: 4.68; p = 0.002). The main effect of session was not significant (est: −0.30 ppm; CI: −0.76-0.14; SE: 0.24; p = 0.212). The group by session interaction was not significant (est: 0.11 ppm; CI: −0.55-0.87; SE: 0.34; p = 0.749).

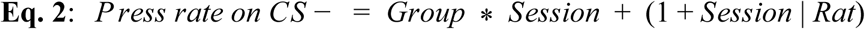

#### Q3. Is CS+ vs. CS-(exp)?

Experimental animals did not display a preference for the fully reinforced lever compared to the partially reinforced lever. A linear mixed model of press rates by lever type and session with random effects of rat (addition of random slopes led to failed convergence; Eq. 3) revealed a non-significant main effect of lever type (Fig. 2C; est: −2.90 ppm; CI: −9.53-3.54; SE: 3.14; p = 0.357). There was not a significant main effect of day (est: −0.19 ppm; CI: −0.76-0.40; SE: 0.30; p = 0.525) nor day by lever type interaction (est: 0.44 ppm; CI: −0.38-1.38; SE: 0.43; p = 0.270).

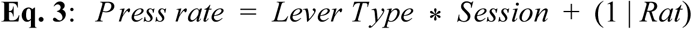

#### Q4. Is CS+ vs. CS-(cont)?

Control animals were able to discriminate between the fully reinforced CS+ and never reinforced CS-. A linear mixed model of press rates by lever type and session with random effects of slope and rat (Eq. 4) revealed a significant main effect of lever type (Fig. 2D; est: 17.4 ppm; CI:10.8-24.1; SE: 3.29; p < 0.001). The main effect of session was not significant (est: −0.30 ppm; CI: −1.09-0.42; SE: 0.38; p = 0.436). The interaction of session by lever type was also not significant (est: 0.15 ppm; CI: −0.73-1.10; SE: 0.45; p = 0.745).

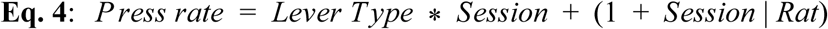

#### Q5. Does magazine entry rate differ between groups?

Animals in the partially reinforced condition did not differ from the fully reinforced condition in magazine entry rates (mpm; magazine entries per minute (total magazine entries/60 min session)). A linear mixed model of magazine entry rates by group and session with random effects of slope and intercept (Eq. 5) revealed a significant main effect of session (Fig. 2E; est: −0.49 mpm; CI: −0.68-(−0.28); SE: 0.11; p < 0.001). The main effect of group was not significant (est: 1.96 mpm; CI: −1.97-6.12; SE: 2.16; p = 0.372). The interaction between session and group was not significant (est: 0.29 mpm; CI: 0.01-0.57; SE: 0.15; p = 0.053)

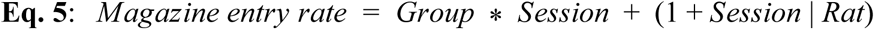

### Extinction

#### Q1. Is CS+ (exp) different from CS+ (control) in extinction?

Both groups decreased pressing on the fully reinforced CS+ over the four days of extinction. A linear mixed model of press rates by session and group with random effects of slope and rat (Eq.6) revealed a significant main effect of session (Fig. 3A; est: −4.64 ppm; CI: −7.26-(−1.92); SE: 1.31; p = 0.003). Groups did not differ in press rates (est: 1.64 ppm; CI: −11.2-14.5; SE: 6.51; p = 0.805). The interaction of group by session was not significant (est: −0.753 ppm; CI: −4.45-3.13; SE: 1.86; p = 0.690).

**Figure 3.**
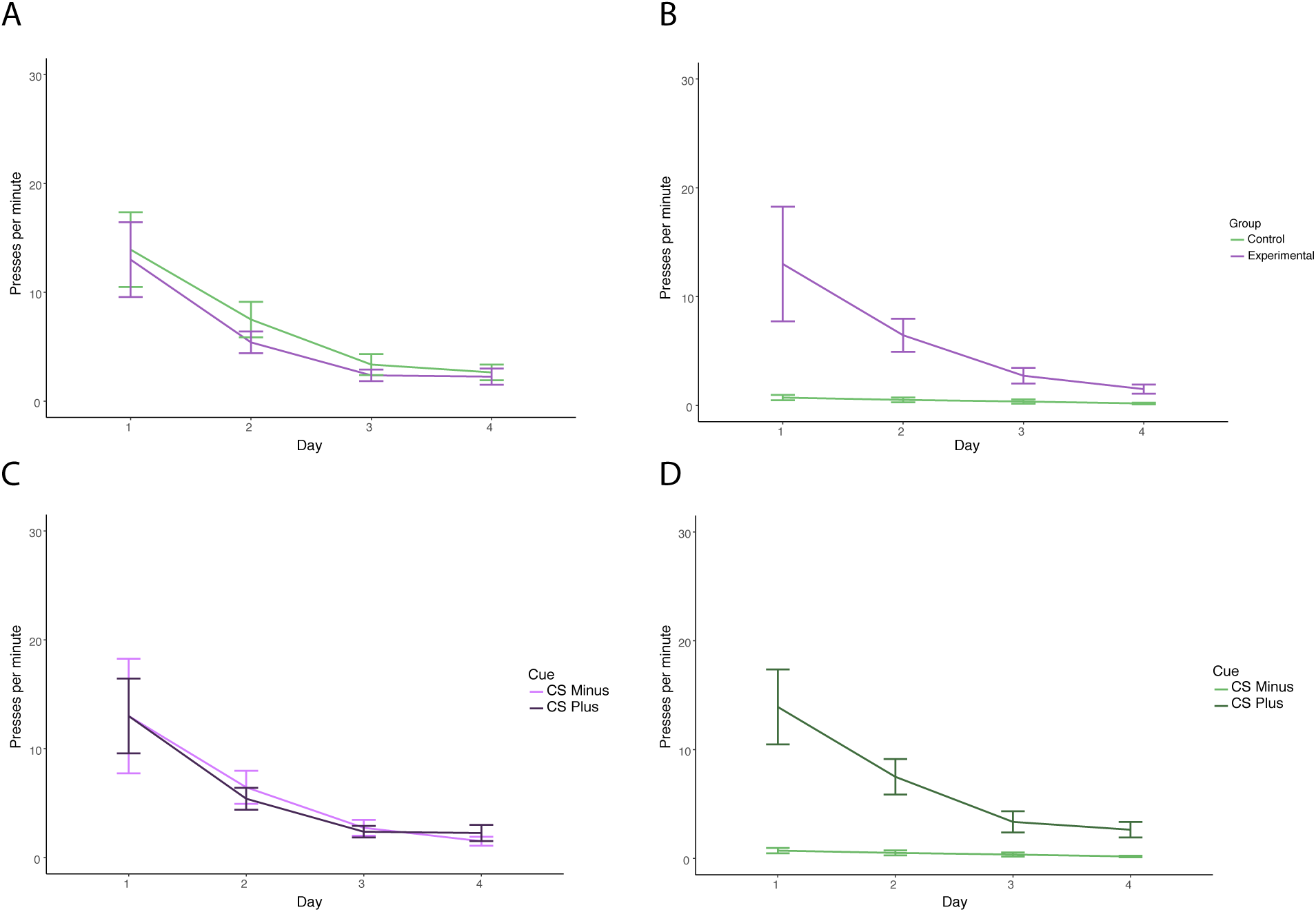
Experiment 1: Extinction. Press rates from the 4 extinction days following training. All error bars represent standard error of the mean. (A) Presses per minute on CS+ during extinction. Group Control (Control; 100%; green) Group 2 (Experimental; 100%; pink). (B) Presses per minute on CS-during extinction. Group 1 (Control; 0%; green) Group 2 (Experimental; 50%; pink). (C) Presses per minute on the CS-(50% reinforced; light pink) and CS+ (100% reinforced; dark pink). (D) Presses per minute on the CS- (0% reinforced, light green) and the CS+ (100% reinforced, dark green).

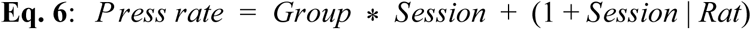

#### Q2. Is CS-(partially reinforced; exp) different from CS-(never reinforced; control) in extinction?

Experimental animals took longer to extinguish the partially reinforced CS compared to control animals. A linear mixed model of press rates by group and session with random effects of slope and rat (Eq. 6) revealed a significant main effect of group (Fig. 3B; est: 21.6 ppm; CI: 10.0-33.7;SE: 6.19; p = 0.003). The main effect of session was not significant (est: −0.18 ppm; CI: −2.31-2.33; SE: 1.26; p = 0.890) however, the interaction of session by group was significant (est: −5.75 ppm; CI: −9.30-(−2.35); SE: 1.79; p = 0.005).

#### Q3. Is CS+ vs. CS-(exp) in extinction?

Both the CS+ and CS-levers extinguish at the same rate for the experimental animals. A linear mixed model of press rates by lever type and session with random effects of slope and rat (Eq. 4) revealed a significant main effect of session (Fig. 3C; est: −5.93 ppm; CI: −9.21-(−3.13); SE: 1.60; p = 0.003). Lever type did not differ as indicated by an non-significant main effect of lever type (est: −1.68 ppm; CI: −8.72-5.11; SE: 3.36; p = 0.619). Press rates to either lever did not differ over extinction sessions (est: 0.53 ppm; CI: −1.79-2.90; SE: 1.23; p = 0.669).

#### Q4. Is CS+ vs. CS-(cont) in extinction?

The control animals display an extinction curve for the CS+ that decreases over sessions while maintaining a near zero press rate towards the CS-. A linear mixed model of press rates by lever type and session with random effects of slope and rat (Eq. 4) revealed a main effect of lever type, with greater overall pressing towards the CS+ (Fig. 3D; est: 18.3 ppm; CI: 13.9-22.6; SE: 2.34; p < 0.001). Pressing towards the CS+ decreased over extinction sessions as seen in a significant interaction of lever type by session (est: −4.47 ppm; CI: −6.07-(−2.78); SE: 0.85; p < 0.001) however, the main effect of session was not significant (est: −0.18 ppm; CI: −1.65-1.45; SE: 0.761; p = 0.819).

### Experiment 2: Long CS Training

Only 7 animals are reported, one animal displayed highly aggressive behavior on day 9 of training and was not run for the remainder of the experiment. Data reflects the absence of this animal. Further, an oversight led to animals trained on a standard (10s CS+, 10s CS-program) on day 1 of training. This day was removed from graphing and analysis (such that there are 11 days of training data displayed and analyzed).

#### Q1. Can animals discriminate between long CS+ vs. CS-levers?

Within animals comparisons of CS+ to CS-demonstrated the animal’s ability to discriminate between rewarded and non-rewarded long cues. A linear mixed model (Eq. 4) showed a significant main effect of lever type with the rewarded lever having greater lever pressing on average than the non-rewarded lever overall (Fig. 4A; est: 42.8 ppm; CI: 29.6-58.2; SE: 7.43; p < 0.001). Overall, press rates did not differ over sessions as indicated by a non-significant main effect of session (est: 0.31 ppm; CI: −0.80-1.27; SE: 0.55; p = 0.570). Preferences for the CS+ lever developed quickly and maintained as indicated by a non-significant lever type by session interaction (est: −0.88 ppm; CI: −3.14-1.12; SE: 1.10; p = 0.421).

**Figure 4.**
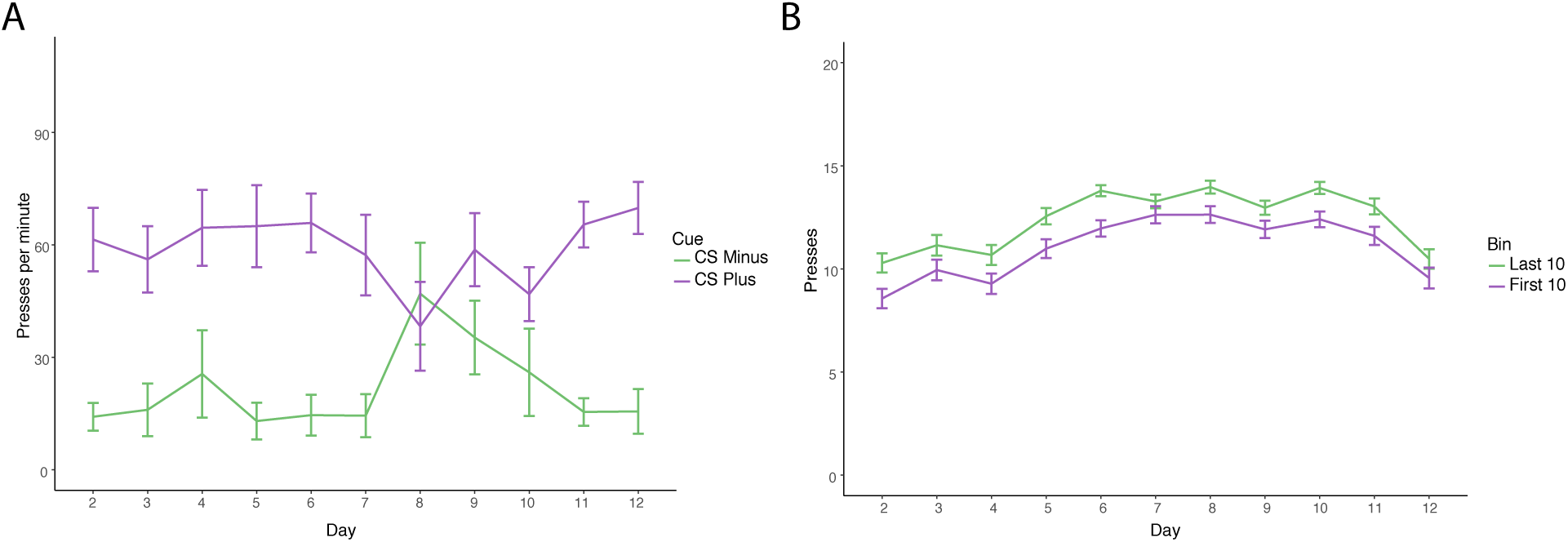
Experiment 2: Training. Training data from the first 11 days of sign-tracking in the Long CS experiment. Error bars are standard error of the mean. (A) Presses per minute on the CS+ and CS-long levers. Presses per minute on the CS+ (purple) and CS-(green) are reported for training days 2 −11. Error bars represent standard error of the mean. (B) Average presses in the first and last 10 seconds of the CS+. Average number of presses in the first (above, green) and last (below, purple) ten seconds of the CS+ presentation for training days 2 − 11. Error bars represent standard error of the mean.

#### Q2. Is the first half of the cue press sums different from the second half of the cue?

Within animals, comparisons of the first 10 sec of CS+ to the last 10 sec of CS+ presentation revealed no difference in pressing earlier in the lever presentation as seen in previous reports of serial discrete levers. A linear mixed model analyzed the sum of presses by the bin (first v. last 10 sec) and session with random effects of session and animal (Eq. 7). Analysis revealed a main effect of bin with a difference between the first and last ten seconds of the cue with the earlier part of the cue (the first ten seconds) having fewer presses (Fig. 4B; est: −1.56 presses; CI: −2.21-(−0.93); SE: 0.35; p < 0.001). Session was not significant (est: 0.18 presses; CI: −0.02-0.35; SE: 0.10; p = 0.117) nor the session by bin interaction (est: 0.04 presses; CI: −0.06-0.14; SE: 0.05; p = 0.460).

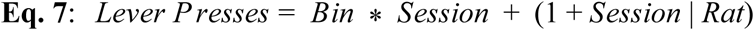

### Extinction

#### Q1. Compare extinction of a short lever (10s - group control) versus long lever (first 10s of 20s lever - Long CS Group)

Comparing rates of extinction on the CS+ between animals who received only CS+ lever extinction (Group Long CS) versus animals who received both levers in extinction (Group Control) revealed that groups do not differ in press rates during the 10 seconds of lever availability in extinction. A linear mixed model analyzing presses during the first 10 seconds of lever availability by group and day with random effects of slope and rat (Eq. 8) showed no significant main effect of group (Fig. 5A; est: −3.21 presses; CI: −6.50-0.10; SE: 1.78; p = 0.115). Session was significant (est: −1.27 presses; CI: −1.78-(−0.77); SE: 0.27; p = 0.002) indicating a decrease in presses by the end of training. The interaction of group by session was not significant (est: 0.74 presses; CI: −0.02-1.49; SE: 0.41; p = 0.114) as both groups decreased pressing similarly over extinction.

**Figure 5.**
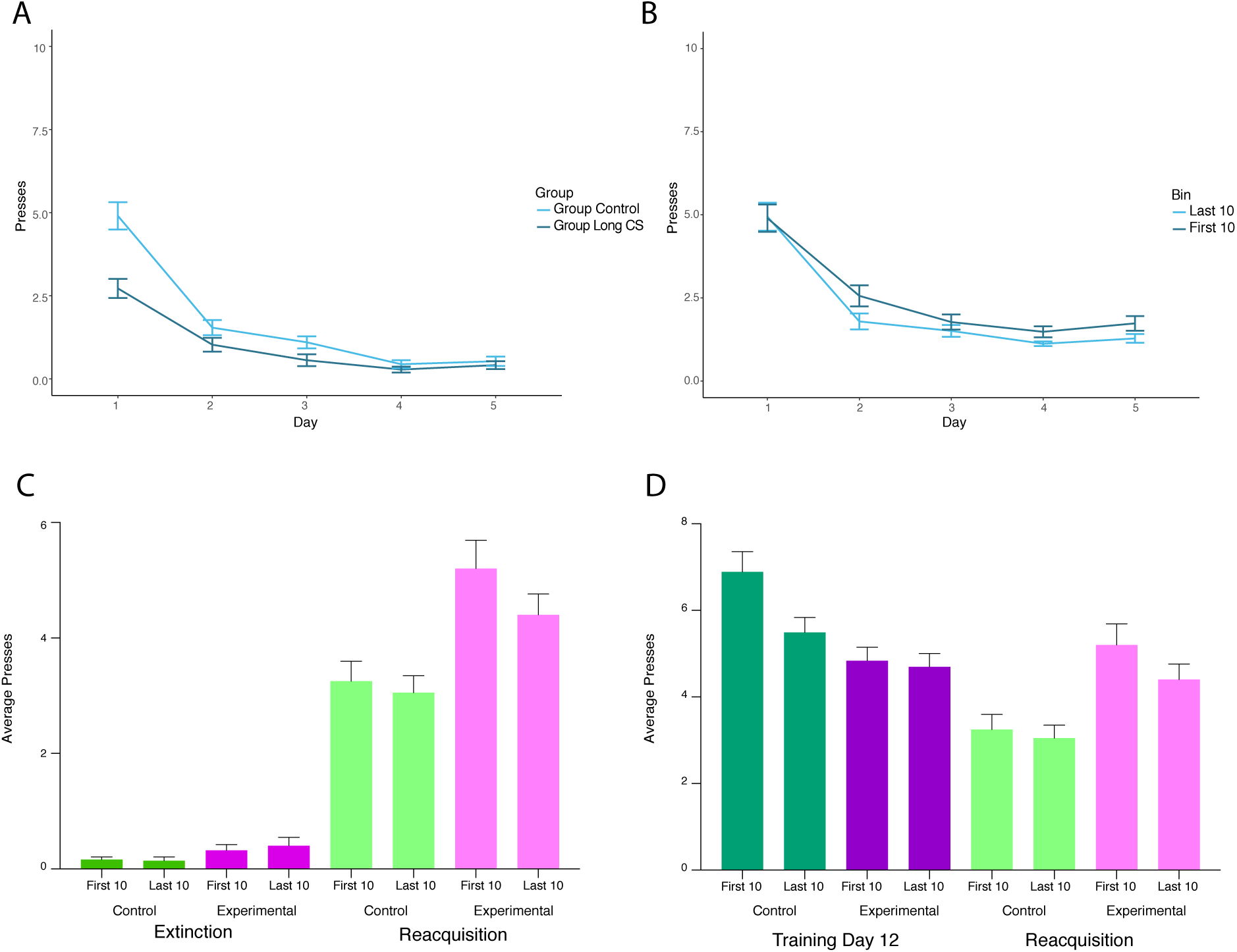
Experiment 2: Extinction and testing. Extinction and test phases of the long CS experiment. Error bars are standard error of the mean. (A) Average presses in the first ten seconds toward the CS in extinction in the control (light blue) and experimental (dark blue) groups. (B) Average presses in the first and last 10 seconds of the CS in the experimental group only during extinction (first 10 seconds, dark blue; last 10 seconds, light blue). (C) Average presses during the Extinction test and reacquisition test in the control (pink) and experimental (green) groups. The first and last 10 second bins are plotted adjacent to each other within groups by test day. (D) Average pressing during the last day of testing (darker colors) and the reacquisition test (lighter colors). The first and last 10 second bins are plotted adjacent to each other within groups by session.

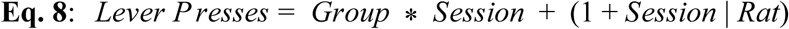

#### Q2. Compare within experimental longCS animals if extinction happens early on in lever or later

Within the long CS group, animals did not differ in extinction rates between early and late lever pressing such that lever presses seemed to extinguish evenly across early and late lever delivery. A linear mixed model of presses by bin (first 10 v. last 10) by session and random effects of slope and rat (Eq. 7) showed a non-significant main effect of bin (Fig 5B; est: 0.21 presses; CI: −0.60-1.01; SE: 0.42; p = 0.621). Session was significant (est: −0.94 presses; CI: −1.36-(−0.46); SE: 0.24; p = 0.024) indicating a decrease in pressing by the end of training. The interaction of session and bin was not significant (est: 0.05 presses; CI: −0.20-0.31; SE: 0.13; p = 0.695), presses in both early and late bins decreased similarly by the end of extinction in the long CS group.

#### Q3. Extinction test v. reacquisition test: Do groups differ in how they press to early v. late lever depending on if it is rewarded or not?

The Long CS group (who received extinction of a long CS) and the Control group (who received extinction of a short CS) did not differ during extinction nor reacquisition testing in presses toward their CS+. A linear mixed model of presses by a three way interaction of group (LongCS vs. Control) by test day (Extinction Test vs. Reacquisition Test) by bin (first 10 seconds vs. last 10 seconds) with random effects of slope and day (Eq. 9) showed no main effect of group (Fig. 5C; est: −1.63 presses; CI: −5.42-2.36; SE: 1.80; p = 0.391) nor bin (est: 0.16 presses; CI: −0.80-1.31; SE: 0.57; p = 0.781) however the main effect of test day was significant (est: 3.09 presses; CI: 0.77-5.63; SE: 1.19; p = 0.034). Groups did not differ between test days (est: 1.79 presses; CI: −2.23-5.66; SE: 1.82; p = 0.356) nor did pressing in either the early or late bins of the CS+ differed between days (est: −0.18 presses; CI: −0.85-0.41; SE: 0.36; p = 0.620). Groups did not differ by early or late press bins by test day (est: −0.70 presses; CI:-1.82-0.33; SE: 0.55; p = 0.208)

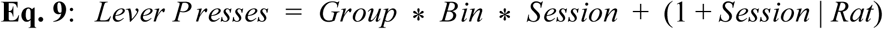

#### Q4. Day 12 of Testing v. Reacquisition test: Do groups reacquire to similar rates seen on Day 12 of training?

Rats did not reacquire to the same level of pressing as seen on the last day of testing. Overall both groups pressed fewer times in the reacquisition test than in testing. Across both sessions, the last 10 seconds of the lever presentation has fewer presses than the first 10 seconds of the cue (i.e. a bias for the first 10 seconds of the cue), this was especially true during the reacquisition test. Further, there is an interaction of the difference between first and last bins by group and day, as well as group by bin. To elaborate, the difference between the first and last bins for control animals on day 12 is greater than the difference on reacquisition day and the opposite is true for the experimental group. A linear mixed model of presses by group by day by bin with random effects of slope and rat (Eq. 9) showed a significant difference of day (Fig. 5D; est: −1.69 presses; CI: −3.03-(−0.27); SE: 0.63; p = 0.011) with the reacquisition day showing fewer presses than day 12. Groups did not differ between days as seen in a non-significant main effect of group (est: 0.17 presses; CI: −4.84-4.44; SE: 1.85; p = 0.928) and non-significant interaction of group by day (est: 0.48 presses; CI: −2.12-3.38; SE: 1.14; p = 0.673). There was a general preference toward the first 10 seconds of the cue (est: −2.60 presses; CI: −4.04-(−1.06); SE: 0.77; p < 0.001) as well as an interaction of bin by day (est: 1.20 presses; CI: 0.85-5.41; SE: 0.49; p = 0.140). Further, groups differed by bin preference (est: 3.11 presses; CI: 0.26-2.10; SE: 1.18; p = 0.008) and by bin preference by day (est: −1.85 presses; CI: −3.36-(−0.48); SE: 0.74; p = 0.013).

### Experiment 3: Backwards CS Training

#### Q1. Do groups differ in CS+ presses?

The backwards CS+ group had a lower rate of pressing towards their backwards CS+ compared to the forward CS+ in the control group. A linear mixed model of press rates on the CS+ by group and session with random effects of slope and rat (Eq. 6) revealed a significant main effect of group (Fig. 6A; est: −23.4 ppm; CI: −30.1-(−16.8); SE: 3.29; p < 0.001). Session was significant (est: 1.46 ppm; CI: 0.88-2.11; SE: 0.30; p < 0.001). The group by session interaction was significant (est: −1.47 ppm; CI: −2.32-(−0.69); SE: 0.42; p = 0.002).

**Figure 6.**
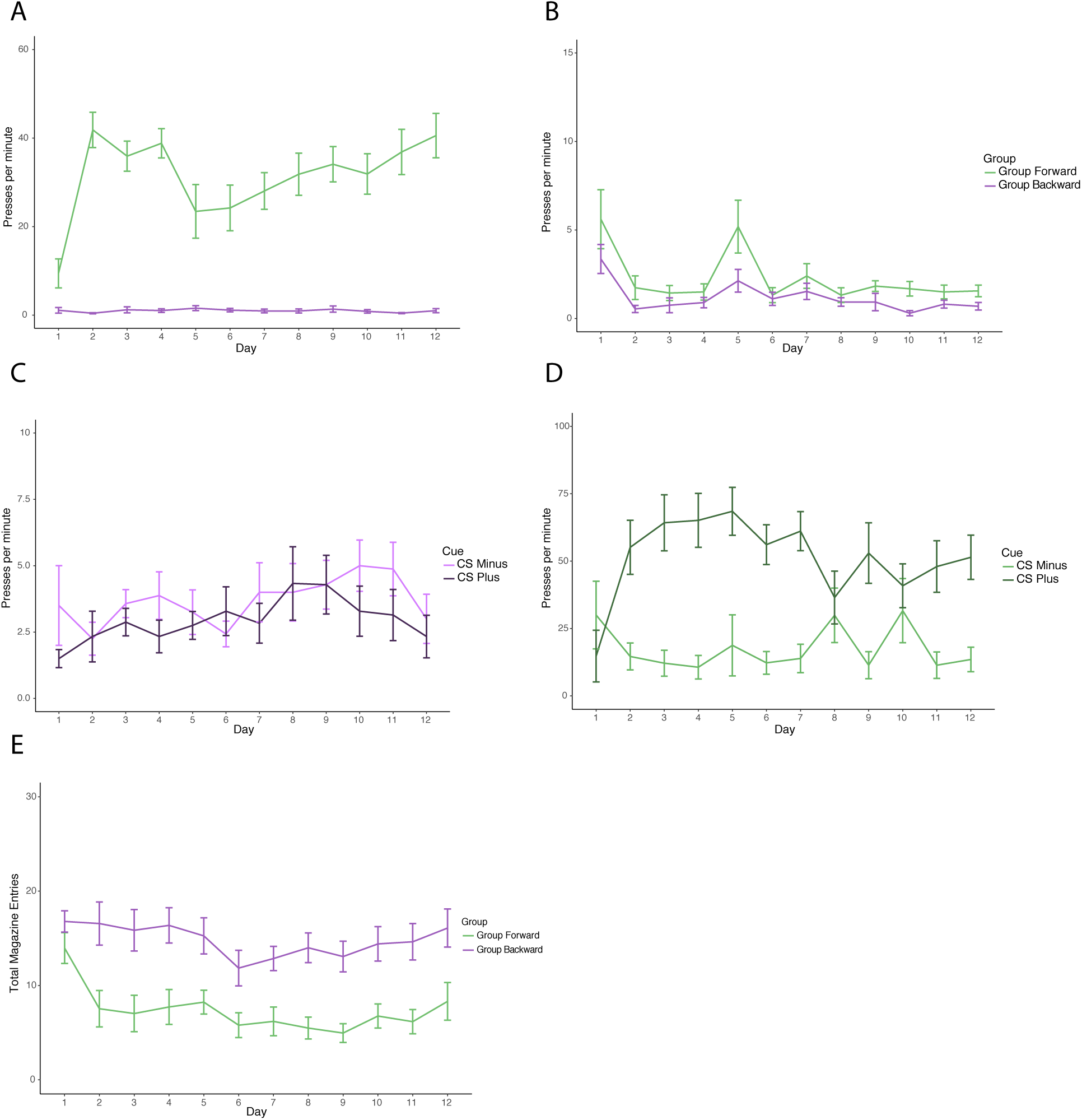
Experiment 3: Training. Press rates from the 12 training days. All error bars represent standard error of the mean. (A) Presses per minute on CS+ during in Group Forward (green) and Group Backward (pink). (B) Presses per minute on CS-in Group 1 Group Forward (green) and Group Backward (pink). (C) Presses per minute on the CS-(light pink) and CS+ (dark pink) in Group Backward. (D) Presses per minute on the CS-(light green) and the CS+ (dark green) in Group Forward. (E) Magazine entries per minute in Group Forward (green) and Group Backward (pink).

#### Q2. Do groups differ in CS-presses?

The backwards CS+ group and the forwards CS+ group did not differ in their rates of pressing towards a CS-. A linear mixed model of press rate on the CS-by group by session with random effects of slope and rat (Eq. 6) identified a non-significant main effect of group (Fig. 6B; est: −1.57 ppm; CI: (−3.22-0.29); SE: 0.88; p = 0.093). Session was significant in that press rates were lower by the end of training (est: −0.19 ppm; CI: −0.33-(−0.05); SE: 0.07; p = 0.011). The group by session interaction was not significant, both groups lowered press rates similarly over sessions (est: 0.07 ppm; CI: −0.12-0.26; SE: 0.10; p = 0.454).

#### Q3. Does backward CS+ differ from CS-in the experimental group?

Animals in the backwards condition were not able to discriminate between the backwards CS+ and the CS-. A linear mixed model of press rates by lever type and session with random effects of slope and rat (Eq. 4) revealed a non-significant main effect of lever type (Fig. 6C; est: −8.83 ppm; CI: −19.3-3.24; SE: 5.30; p = 0.097). Session was not significant (est: −0.77 ppm; CI: −1.75-0.28; SE: 0.51; p 0.135). The session by lever type interaction was not significant (est: 1.13 ppm; CI: −0.45-2.52; SE: 0.72; p = 0.118).

#### Q4. Does CS+ differ from CS-in control group?

Animals in the forward condition were able to discriminate between the CS+ and CS-. A linear mixed model of press rates by lever type and session with random effects of slope and rat (Eq. 4) identified a significant main effect of lever type (Fig. 6D; est: 39.7 ppm; CI: 26.6-53.0; SE: 7.16; p < 0.001). Session was not significant (est: −0.85 ppm; CI: −2.30-0.62; SE: 0.73; p = 0.256). The lever type by session interaction was not significant (est: −0.24 ppm; CI: −2.11-1.55; SE: 0.97; p = 0.802).

#### Q5. Do groups differ in overall magazine entries?

The backwards CS+ group displayed greater overall magazine entries than the forward CS+ group. A linear mixed model of magazine entry rates (mpm; total magazine entries over the session divided by session time (avg. 60 min)) by group and session by random effects of slope and rat (Eq. 5) revealed a significant main effect of group (Fig. 6E; est: 7.97 mpm; CI: 3.07-13.1; SE: 2.62; p = 0.007). Session was also significant (est: −0.34 mpm; CI: −0.59-(−0.07); SE: 0.13; p = 0.009). The group by session interaction was not significant (est: 0.19 mpm; CI: −0.15-0.53; SE: 0.18; p = 0.298).

### Reversal and Reversal Test - Lever Responding

#### Q1. Does Group Forward differ from Group Backward in press rates toward CS+?

Group Backward displayed greater pressing towards their now forward CS+ than Group Forward toward their now backward CS+ in the reversal period. A linear mixed model of press rates by group and session with random effects of slope and rat (Eq. 1) showed a main effect of group (Fig. 7A; est: −35.1 ppm; CI: −66.4-(−2.85); SE: 16.4; p = 0.044). The main effect of session was not significant (est: −1.57 ppm; CI: −3.04-(−0.02); SE: 0.75; p = 0.0562) however the group by session interaction was significant (est: 2.37 ppm; CI: 0.41-4.40; SE: 1.07; p = 0.043) with the backward group showing greater press rates toward their now forward CS+ compared to the lessening press rates in the forward group on their now backward CS+.

**Figure 7.**
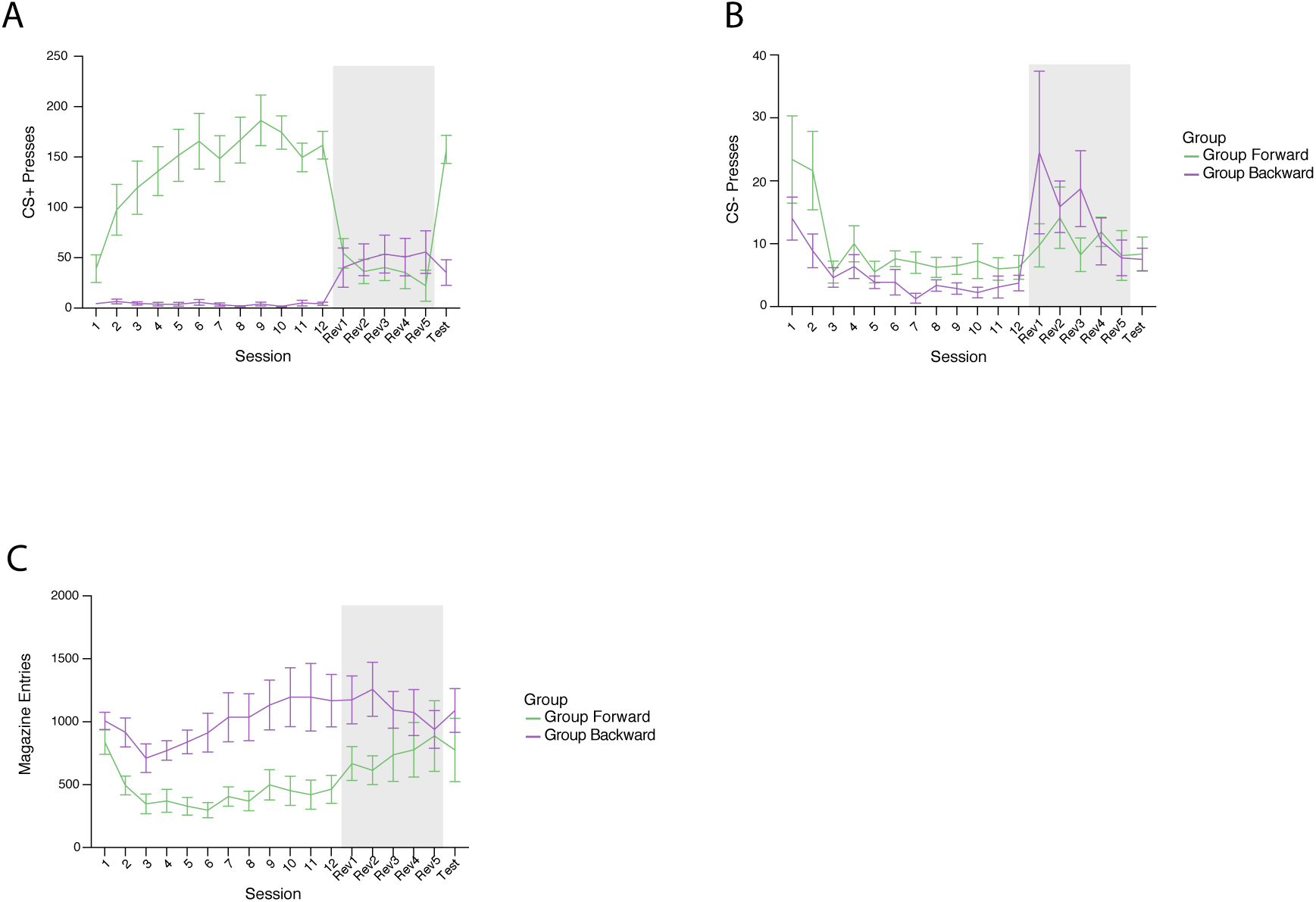
Experiment 3: Reversal and Reversal Test in relation to previous training. Graphs show a summary of training, reversal, and test days to see broad trends in behavior. Grey blocks depict reversal period while pink areas are the reversal test. Error bars represent SEM. (A) CS+ responding (CS+ is assigned as the CS+ lever during training, graphs depict this lever across all phases of the experiment) in forward (black) and backward (green) groups. (B) CS-responding in forward (black) and backward (green) groups. (C) Magazine entries (totals) in the forward (black) and backward (green) groups.

#### Q2. Does Group Forward differ from Group Backward in press rates toward CS-?

Group Backward and Group Forward did not differ in their press rates toward the CS never paired in any capacity with reward (the CS-) in the reversal period. A linear mixed model of press rates by group by session with random effects of slope and rat (Eq. 2) showed no significant difference between groups (Fig. 7B; est: 0.10 ppm; CI: −4.82-4.77; SE: 2.47; p = 0.968). Sessions did not differ (est: −0.14 ppm; CI: −0.36-0.04; SE: 0.11; p = 0.195) nor did groups differ over sessions (est: −0.06 ppm; CI: −0.35-0.25; SE: 0.15; p = 0.685).

#### Q3. Can Group Forward discriminate between CS+ and CS-?

Group Forward was unable to discriminate during the reversal period between the now backward CS+ and the CS-. A linear mixed model of press rates by lever type by session with random effects of slope and rat (Eq. 4) showed a non-significant main effect of lever type (Fig. 7A,B; est: 11.5 ppm; CI: 1.44-20.23; SE: 4.89; p = 0.021). Sessions did not differ (est: −2.02 ppm; CI: −4.25-(−0.01); SE: 1.06; p = 0.062) nor did lever type over sessions (est: 3.74 ppm; CI: 0.91-6.86; SE: 1.47; p = 0.013).

#### Q4. Can Group Backward discriminate between CS+ and CS-?

Group Backward was unable to discriminate between the now forward CS+ and CS-cues in the reversal program. A linear mixed model of press rates by lever type by session with random effects of slope and rat (Eq. 4) revealed a non-significant difference between the CS+ and CS-levers (Fig. 7A,B; est: −1.48 ppm; CI: −10.4-8.17; SE: 4.63; p = 0.751). Press rates did not differ by day (est: −1.39 ppm; CI: −3.52-0.75; SE: 1.03; p = 0.185) nor was there a significant difference between levers over reversal sessions (est: 0.70 ppm; CI: −2.14-3.20; SE: 1.40; p = 0.617).

### Reversal Test - Lever Responding

#### Q5. Do rates in fwd group differ in CS+ and CS-on test day?

Forward Group animals were able to discriminate between the CS+ and CS-on the reversal test day. A paired t-test of presses on test day by lever type in the forward group revealed a significant difference in lever presses with greater pressing toward the CS+ on the reversal day (t(39) = −9.046; CI: −27.8-(−17.6); p < 0.001).

#### Q6. Do rates in bkwd group differ in CS+ and CS-on test day?

Backward Group animals were unable to discriminate between the CS+ and CS-on the reversal test day. A paired t-test of presses on the test day by lever type in the backward group revealed a non-significant difference in lever presses (t(39) = −0.282; CI: −5.11-3.86; p = 0.780).

### Reversal and Reversal Test - Magazine Entries

#### Q1. Does Group Forward differ from Group Backward in Magazine entry rates in Reversal period?

The Forward Group and the Backward Group did not differ in magazine entry rates during the reversal period. A linear mixed model of magazine entry rates by group and session with random effects of slope and rat (Eq. 5) showed no significant differences between groups (Fig. 7C; est: −6.88 mpm; CI: −23.8-9.55; SE: 8.67; p = 0.439) nor differences by session (est: 0.12 mpm; CI: −0.74-0.97; SE: 0.47; p = 0.798). Groups do not differ over sessions (est: 1.09 mpm; CI: −0.16-2.40; SE: 0.66; p = 0.120).

#### Q2. Does Group Forward differ from Group Backward in Magazine entry rates in Reversal Test?

The Forward Group and the Backward Group differed in magazine entry rates during the reversal test. An unpaired, Welch two sample t-test of magazine entries per minute by group showed significant differences in magazine entries by group (t(11.5) = −2.745; CI: −19.0-(−2.14); p = 0.018).

## Discussion

Experiment one demonstrated that manipulation of the reward structure of one lever cue does not impact the responding towards a fully reinforced lever. Control and experimental animals pressed equally to the fully reinforced lever, even though group experimental pressed the partially reinforced lever more than group control pressed the never reinforced lever. Additionally, group experimental did not display a preference for the fully reinforced lever over the partially reinforced lever and extinguished the two levers at the same rate.

These results do not align with prior findings that partial reinforcement of the conditioned stimulus results in more sign-tracking behavior while full reinforcement leads to more frequent goal-tracking behavior (Davey and Cleland, 2013). Additionally, this experiment contradicts research that found that responding declined more slowly to the partially reinforced stimulus compared to the fully reinforced stimulus during extinction (Davey and Cleland, 2013).

When animals were given 20sec CS+ and 20sec CS-levers in experiment two, they were able to discriminate between the two. Results also showed that animals had similar press rates in the first 10 seconds of the CS+ as they did during the last 10 seconds of the CS+. In previous studies where a sequential presentation of lever cues occurred, the lever presented furthest in time from the reward delivery acquired more lever responding than the lever temporally closer in time to the reward over sessions (Smedley and Smith 2018a, 2018b). However, the results of experiment two suggest that this phenomenon may be unique to discrete lever presentations. Both the Long CS group and the control group, which extinguished with a 10 second CS+, extinguished at similar rates. When both groups returned to the training paradigm in extinction testing, the two groups maintained low rates of responding. Responding increased in both groups when the training was rewarded once again, but responding was less than it had been on the final days of training. This experiment had limitations such as a low number of subjects (n=7), 11 days of training, and analysis of only lever deflections as a measure of responding, rather than including other conditioned responses such as sniffing, licking, and magazine behavior.

The results of the third backwards conditioning experiment showed that animals do not sign-track towards the backwards conditioned stimulus as they do toward the forwards conditioned stimulus. While animals in the forward CS+ group were able to discriminate between the forwards CS+ and CS-, animals in the backwards condition were not able to discriminate between the backwards CS+ and the CS-. The two groups also differed during training in that the backwards CS+ group displayed greater overall magazine entries. This may suggest that the backwards CS+ did not act as an inhibitor, signaling the start of the intertrial period, as mentioned in prior experiments (Chang, Blaisdell, & Miller, 2003). Future research could alter this experiment to compound the backwards CS with a novel tone or light, to assess if the backwards CS had conditioned inhibitory properties.

When the groups in experiment three received a reversal to the reward structure they were given in training, Group Backward displayed greater pressing towards their now forward CS+ than Group Forward displayed toward their now backward CS+. The results also showed that neither group was able to discriminate between their new CS+ and the CS-during the reversal period or on the reversal test day. Unlike during training, the Group Forward and Group Backward did not differ in magazine entry rates during the reversal period, but Group Backward did have more magazine entries on the reversal test day.

Together, these experiments provide evidence that manipulation of the reinforcement of a conditioned stimulus, presentation of a longer conditioned stimulus instead of two discrete conditioned stimuli, and reversal of the presentation sequence of the conditioned stimulus and the reward can alter sign-tracking behavior.

